# Direct Force Measurement and Loading on Developing Tissues in Intact Avian Embryos

**DOI:** 10.1101/2022.06.20.496880

**Authors:** Chon U Chan, Fengzhu Xiong, Arthur Michaut, Joana M. N. Vidigueira, Olivier Pourquie, L. Mahadevan

## Abstract

Developmental morphogenesis is driven by tissue stresses acting on tissue rheology. Direct measurements of forces in small tissues (100μm-1mm) *in situ* such as in early embryos require high spatial precision and minimal invasiveness. Here we report tissue force microscopy (TiFM) integrating a vertical cantilever probe and live imaging to enable close-loop control of mechanical loading in early chicken embryos. By testing previously qualitatively characterized force-producing tissues in the elongating body axis, we show that TiFM quantitatively captures stress dynamics with high sensitivity. TiFM also provides the capacity of applying a stable, minimally-invasive and physiologically relevant load to drive tissue deformation, which alters morphogenetic progression and cell movements. Together, TiFM addresses a key technological gap in tissue force measurement and manipulation in small developing embryos, and promises to contribute to the quantitative understanding of complex multi-tissue mechanics during development.

## Introduction

During the development of a multicellular organism, cell behaviors collectively generate tissue forces and alter tissue mechanical properties. These changes drive tissue deformation towards functional patterns and shapes. Understanding the dynamics and regulation of these mechanical factors is essential for creating accurate models and controls of tissue morphogenesis in both basic and applied fields of developmental biology, for example organoid engineering. Tissue size remains a major constraint for mechanical studies of early animal embryos, where the fundamental body plan and a variety of distinctly structured and shaped tissues form rapidly at the small scale of 100μm (Mongera *et al*., 2019). At these developmental stages, tissues are very soft and produce small stresses. Currently available *in vivo* approaches suitable for these early embryos include classic embryology methods such as surgical cutting (Schoenwolf and Smith, 1990), cantilever beams and fibres (Hara *et al*., 2013; Chevalier *et al*., 2016), embedding gels (Zhou, Kim and Davidson, 2009), laser ablation (Hutson *et al*., 2009) and emerging (in the sense that they are more recently incorporated for embryos) techniques incorporating precision engineering methods such as magnetic droplets (Serwane *et al*., 2017), atomic force microscopy (AFM) (Barriga *et al*., 2018) and related microindenters (Marrese *et al*., 2020), Brillouin microscopy (Prevedel *et al*., 2019) and optically trapped nanoparticles (Dzementsei *et al*., 2018) among others (Campàs, 2016). These methods provide access to tissue mechanical properties at various resolution and coverage in intact embryos (or large explants), but have limited success in measuring bulk tissue stresses *in situ*.

Using embedded soft alginate gels, we previously detected a pushing force from the axial tissues (neural tube and the notochord) of early chicken embryos (HH8-12, (Hamburger and Hamilton, 1951)) that drives body elongation and cell movement near the posterior progenitor domain (Xiong *et al*., 2020). This pushing force was estimated to be quite small as only very soft alginate gels show marked deformation. The gels were not suitable for accurate quantification of the force as they were heterogenous, irregularly deformed and might undergo mechanical property changes in the chemical environment of the developing embryo. Another general issue with large-size (several to dozens of cell diameters) embedded sensors/actuators is that they cause a large deformation at the local embedding site which could alter the cell organization and tissue mechanics of the normal tissue environment. One way to minimize the tissue impact of force sensors is to use ultra-thin, retrievable probes, which reduces the size and duration of contact required for the measurements. Here we present a new system taking a cantilever deflection approach(Hara *et al*., 2013), which utilizes a beam/needle that is bent when one end is held still and the other end is under a load. By combining modern cantilevers, live imaging and tracking, and electronic sensing in a programmed feedback loop, we constructed a system capable of dynamic force measurement and loading in live avian embryos. In this paper we present the design and validation results of the system, which we termed Tissue Force Microscopy (TiFM), and discuss the considerations in its applications.

## Results and Discussion

The key of accuracy for a cantilever is the precision of the deflection measurement, which comes from the positional difference between the holding end and the loaded end. The bigger the deflection, the better the signal-to-noise ratio. To match the sensitivity required for the small stresses produced by soft body axis tissues in the early chicken embryo, we used commercially available atomic force microscope (AFM) silicon-nitrate probes as our cantilevers. These probes can have low force constants to the order of 0.01N/m (10nN/μm to put in the small tissue perspective). In contrast to the tapping mechanism in AFM surface imaging, we position these thin (~1μm) cantilevers vertically to allow direct insertion into the tissue, with or without modifications to the tip. In the case of measuring the axial pushing force, because the tissue cross-section is much larger than that of the cantilever tip, we glued a tailored piece of aluminium foil (200μm square, ~15μm thick) to the tip (Michaut, 2018), which fully blocks the elongating neural tube and notochord in a HH11 chicken embryo upon insertion. The embryo (prepared using the *ex ovo* EC culture protocol on a piece of windowed filter paper(Chapman *et al*., 2001)) is mounted on a glass bottom dish and imaged from below (Figs. 1A, S1). The glass bottom dish contains a thin layer of culture gel to support the embryo, and the embryo is covered by a thin layer of Phosphate-buffered saline (PBS) and mineral oil to prevent drying (Michaut, 2018). The whole stage is set in an environmental enclosure maintaining 37.5°C with heating fans. Embryos develop normally at a slightly lower rate for at least 6 hours under these conditions as assessed by somite formation and axis elongation (~2hrs per somite as opposed to the ~1.5hrs normal rate, ~100μm/hr elongation speed as opposed to the ~150μm/hr normal rate). Notably, the probe insertion site heals quickly after cantilever retraction and becomes barely distinguishable in a few minutes. The inverted microscope (10x objective) captures images of the tissue section where the probe tip/attached foil is in focus and sends them to the computer for real-time segmentation to measure the position of the tip.

**Figure 1.**
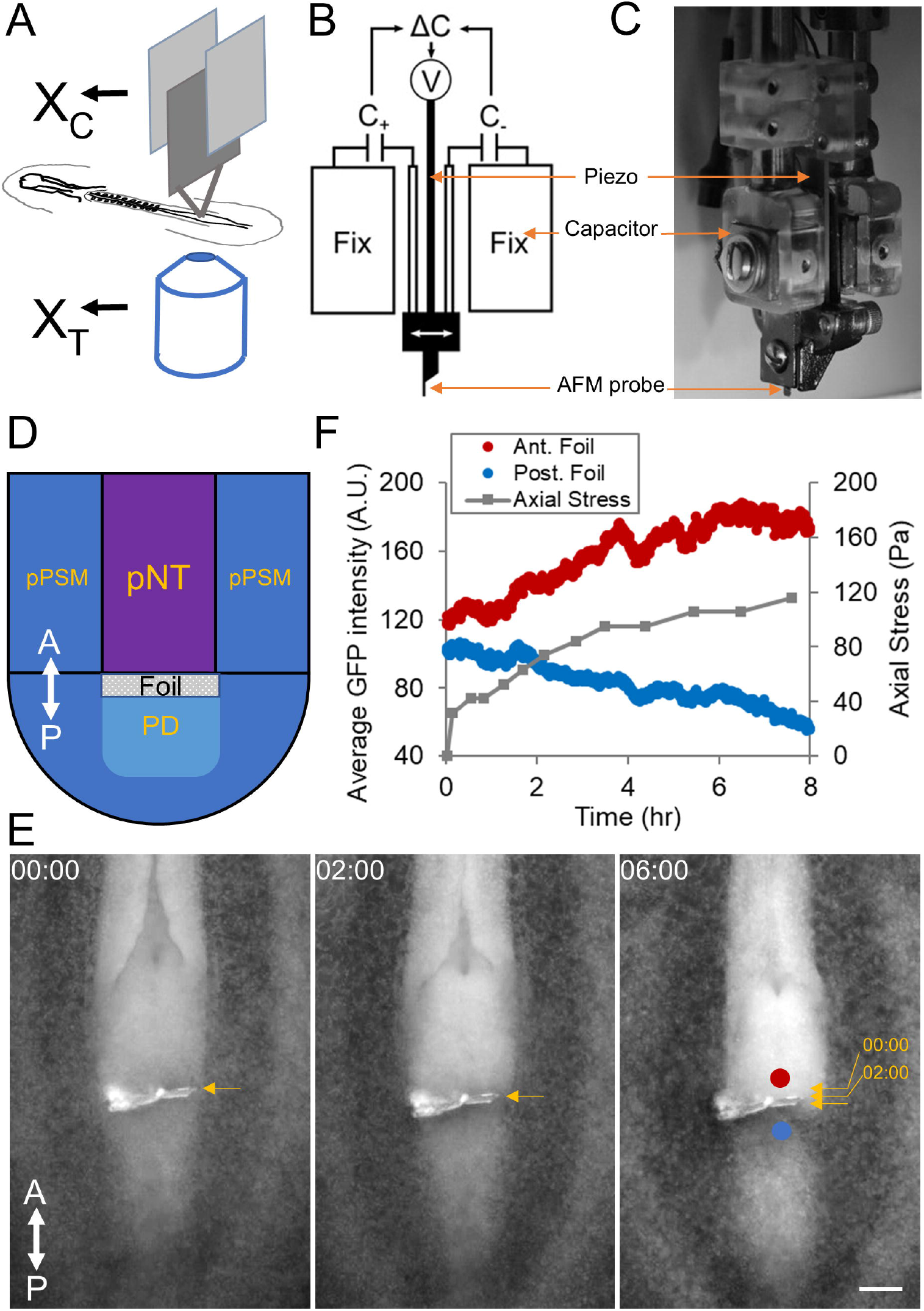
Tissue force microscope (TiFM) to measure the axial elongation force. (A) Concept of TiFM. The design takes advantage of the flatness of the early avian embryo. X_C_ is the holder “chip” position measured by the capacitors, X_T_ is the probe “tip” position measured by the microscope. Their difference measures the deflection of the cantilever beam. (B-C) Probe holder and capacitors. Two capacitor plates and the piezo are integrated for position control and measurement against two fixed plates. C, capacitance; V, voltage; (C) is a side photo of the assembled probe holder. (D) Diagram of the axial elongation stress measurement. This is a dorsal view of the tail end of the embryo as seen in (E), the probe enters from the ventral side. A-P double-sided arrows mark the antero-posterior axis, same for all following Figures where relevant. pPSM, posterior presomitic mesoderm; pNT, posterior neural tube; PD, progenitor domain. Cells from the PD enter the U-shaped PSM under the pushing forces from the pNT and notochord (Xiong *et al*., 2020). (E) Foil movement under the elongation force. Arrows indicate the small displacements of the foil (overlaid on the third image). The foil depth is ~200μm. Blue and Red dots mark the ROIs for fluorescence intensity measurements. This is a GFP embryo. Neural tube folds can be seen to be closing and narrowing. Representative of 5 similar experiments. (F) Axial elongation stress and cell density approximated by fluorescence intensity. The stress is calculated by the displacement, the cantilever spring constant (0.2N/m) and the cross-sectional area of the foil. Scale bar: 100μm.

To enable dynamic positioning of the cantilever, the chip holding the cantilevers is mounted on an electric piezo (Fig. S2). To enable precise measurement of the chip/piezo position, they are further flanked with a pair of capacitors (Figs. 1B-C). The capacitance difference between the pair is highly sensitive to the distance between the capacitor plates therefore the movement of the piezo. Before loading the embryo, the chip position and the capacitance reading are first calibrated with the microscope to create a lookup function where capacitance difference is interpreted as chip position. This real time position information can feedback to the voltage controller connected to the piezo as a closed loop system (Figs. 1B-D, methods). Voltage can thus be adjusted automatically if any drift of the piezo/chip is detected ensuring the stability of chip position. Extended imaging of the chip confirms that the feedback loop maintains stable chip positioning.

By taking the position differences between the foil (measured by the microscope) and the chip (measured by the capacitors) over time, and multiplying the cantilever spring constant (0.2N/m) and dividing by the foil cross-sectional area, we found the initial stress (shortly [under 30min] after probe insertion) to be ~40Pa for axial elongation and the stalling stress in the longer term (>5hrs) to be ~100Pa (Figs. 1D-F). Cells are observed to accumulate anterior to the foil as the foil moves and eventually stalls (Movie S1). Posterior to the foil the cell density markedly reduces, analogous to the effect of a water dam cutting off flow (Fig. 1F). Foil alone directly connected to the holding chip does not show any stable displacement (Movie S2, Fig. S3A). These results are consistent with our previous gel deformation experiments to detect the axis elongation force (Xiong *et al*., 2020). To assess the invasiveness of the foil, which causes significantly larger area of tissue damage than sharp AFM probes (100-400μm wide and ~15μm thick, compared to 30-50μm wide and ~1μm thick), we took advantage of the piezo driver to cut different-sized foil insertion wounds at the body axis end and followed them over time (Fig. S3B). Thin slits (<20μm) that are typically left after stress measurements heal quickly under 1hr, suggesting minimal cellular changes at the wound site (such as epithelialization of the exposed cells which builds tension that prevents wound closure, a common occurrence in tissue microsurgeries that cause mechanical artifacts) and minimal long-term effects on the tissue area (in contrast to large embedded sensors).

We next measured the stress produced by the posterior presomitic mesoderm (pPSM) flanking the body axis and known to be drivers of elongation (Bénazéraf *et al*., 2010). In this case the tissue convergence speed (~15μm/hr) is much slower than axis elongation (~150μm/hr). In addition, because the notochord normally also undergoes tissue-autonomous convergence and resists PSM deformation, inserting the probe between the notochord and pPSM does not distinguish the source of the detected force. To ensure that we can test whether pPSM generates a stress, we positioned the probe next to the pPSM after surgically removing a portion of the posterior notochord (Fig. 2A). Unlike the anterior PSM (aPSM) or somites, pPSM tissue is known to undergo expansion and will fill into this opening after surgery (Xiong *et al*., 2020). A small stress in the range of 10-100Pa is detected which gradually dissipates over several hours (Figs. 2B-C). This is consistent with our previous observations that the pPSM compresses on axial tissues and that the compression disappears at the differentiating aPSM level (Xiong *et al*., 2020). Our system thus enables direct quantitative (to the precision of order of magnitude in this particular case) confirmation of the small forces generated by the tissues of the elongating chicken body axis.

**Figure 2.**
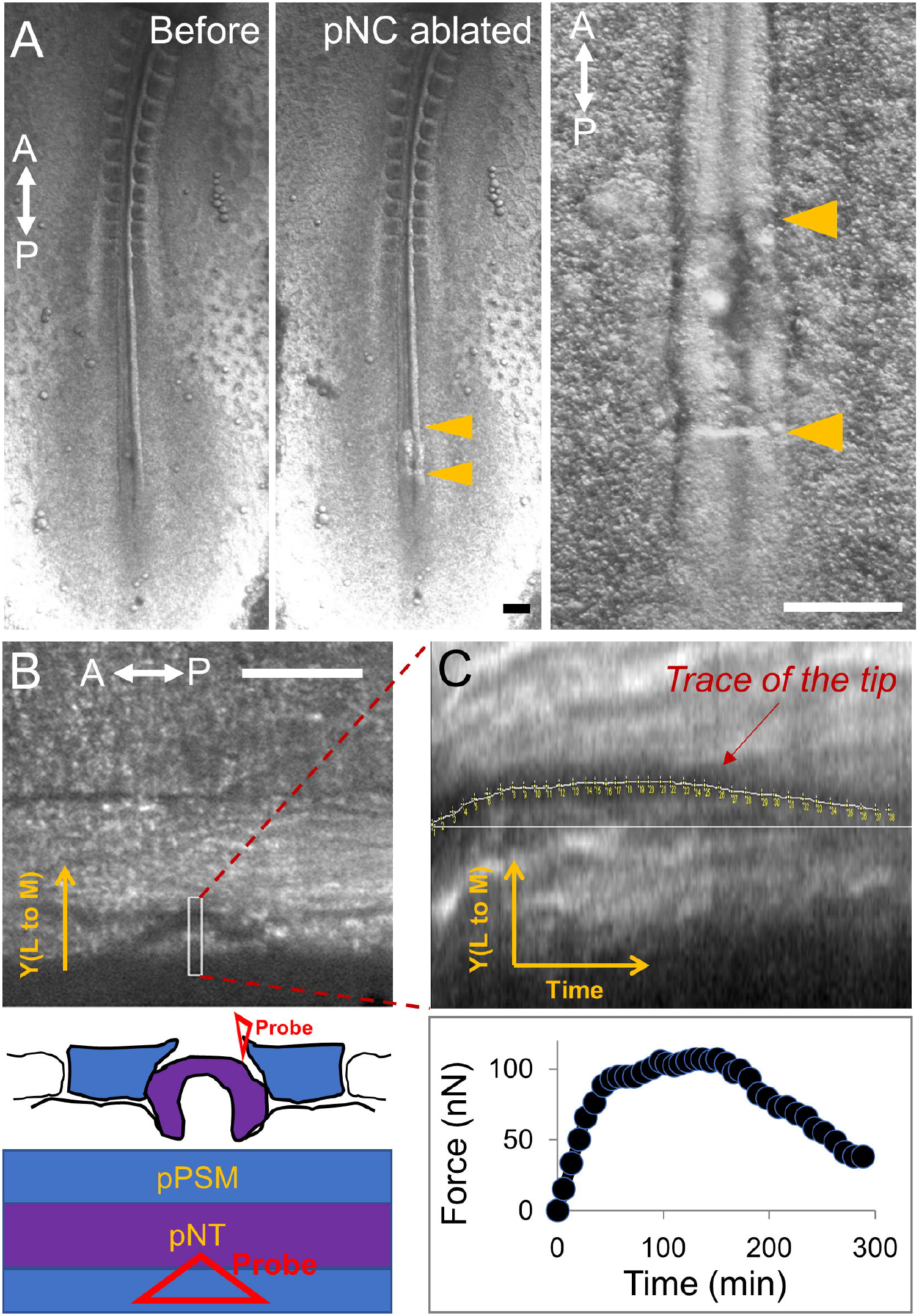
Measurement of the posterior presomitic mesoderm compression. (A) Ventral view of a HH11 embryo undergoing posterior notochord (pNC) ablation to reveal the medial surface of the pPSM. Arrowheads mark the anterior and posterior borders of the surgical window. Neural tube which is further dorsal remains intact and is in the view. Representative of 3 similar experiments. Scale bars: 200μm. (B) Dorsal side view of the region in A now under the TiFM. As indicated in the diagram, a soft triangular probe (0.01N/m) is now inserted medially by the pPSM boundary, the shadow of the triangle tip is visible. The pPSM tissue is known to expand into this area after pNC is ablated. Scale bar: 100μm. (C) Trace of the probe tip shows its deflection over time and translates to the lateral to medial force generated by the pPSM. The force quickly stalls around 100nN and dissipates after a few hours. The estimated depth of the probe is ~30μm and contact surface size with the probe is on the order of 10^3^-10^4^μm^2^, predicting a pPSM stress in the range of 10-100Pa.

To perform controlled mechanical perturbations, we used the feedback loop to move the piezo/chip to maintain a constant deflection by comparing with the tip position obtained with live segmentation of the tip images. This enables a sustained constant force to be applied to the tissue through the cantilever tip. Using this system, we loaded an anterior to posterior steady pulling force (150-200nN) on the axial tissues which at the same time is also a pushing force on the posterior progenitor domain (Fig. 3A). The embryo shows accelerated elongation under this load and surrounding tissues exhibit differently patterned deformations (Figs. 3A-C, Movies S3-4). For example, the neural folds showed clear fastened convergence and closure under loading while PSM elongated without pronounced width change (Fig. 3B). It is also notable that despite being under a strong load that doubled elongation speed (Fig. 3C), the insertion location and surrounding tissues remain intact and no tearing was observed, and the insertion wound quickly disappears after probe retraction (Movie S4). We labelled cell clusters in the pPSM and followed their movements by cell tracking (Fig. 3D). The stress loading causes the cell cluster to move more laterally, following the “U” shaped trajectory from the progenitor domain to the pPSM (Fig. 3E), consistent with more invasive approaches such as a magnetic pin that produces excess stresses beyond the physiological range (Xiong *et al*., 2020). These data together show that in an intact embryo, tissue stresses exerted at one location have wide impacts through inter-tissue connections and alter cell behaviors at a distance, highlighting the importance of integrated multi-tissue models in developmental morphogenesis.

**Figure 3.**
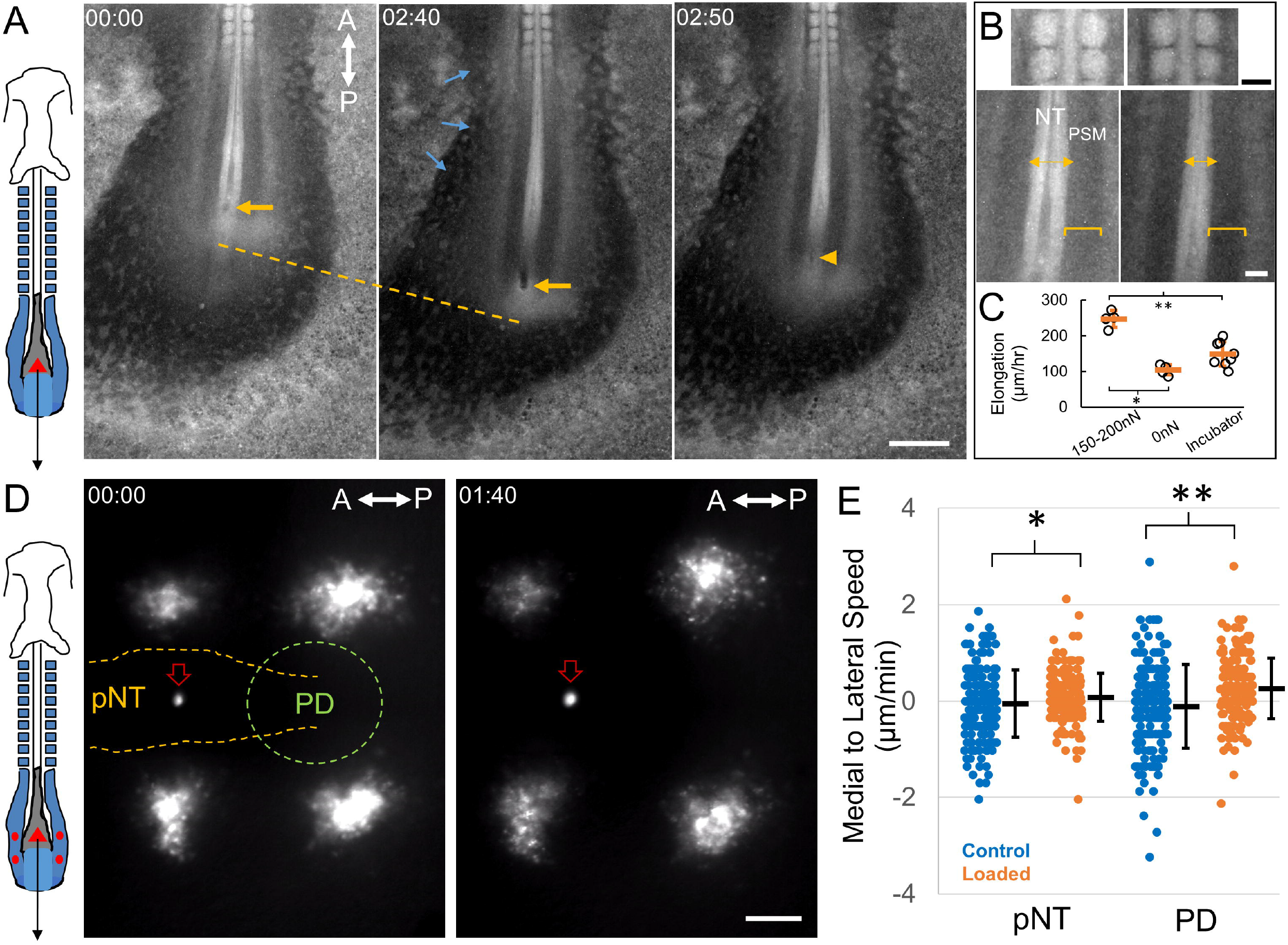
Tissue and cell dynamics following mechanical load. (A) Schematic and images from a loaded embryo. The red triangle on the image indicates the probe tip. The black arrow indicates the direction of the force (same in panel D). Posterior body axis and extraembryonic regions (incl. zona opaca) are visible in the image. This is a GFP embryo. Yellow arrows mark the probe location. Yellow arrowhead shows the wound immediately after probe retraction. Yellow dashed line measures elongation. Blue arrows mark expansion of zona opaca towards the embryo under load. Time stamps are hh:mm. Representative of 5 similar experiments. Scale bar: 500μm. (B) Magnified views of body axis tissues under load, comparing 00:00 and 02:40 timepoints from panel A. Somites and anterior neural tube are largely unchanged. Posterior neural tube markedly narrows while PSM narrows little. NT, neural tube; PSM, pre-somitic mesoderm. Scale bars: 100μm. (C) Physiological loading increases elongation speed. 0nN indicates no load control on the TiFM. Incubator control includes samples not on the TiFM mounting environment, which develop faster. Error bars: mean±s.d.. *,**p<0.001, unpaired 2-tail t-tests. (D) Schematic and images from a loaded and DiI labelled embryo. The probe tip is fluorescent in the red channel similarly as DiI (red empty arrow in the images). Red dots mark the DiI injection sites in the pPSM corresponding to the clusters of cells on the image. pNT, posterior neural tube; PD, progenitor domain. 2 clusters are on the same anterior-posterior level as the pNT and 2 as the PD. Some of the DiI labelled cells in these movies can be tracked to analyse cell movements. Time stamps are hh:mm. Representative of 4 similar experiments. Scale bar: 100μm. (E) Medial to lateral speeds of cells measured from tracks on the pNT and PD levels, respectively. Each speed measurement is taken by the cell’s displacement over a 5-minute interval. Loading causes cells to move more laterally on average. Error bars: mean±s.e.m. *p=0.032; **p<0.001, unpaired 2-tail t-tests.

We term the close-loop electro-mechanical system including the vertical cantilever, the piezo, the capacitors, live imaging and incubation described here as Tissue Force Microscopy (TiFM, Fig. S1). TiFM theoretically reaches a sensitivity of 1nN (limited by the resolution and accuracy of tip imaging and tracking) with the present hardware and has 3D coverage at ~20μm spatial resolution (typical widths of the probe tip) parallel to the stress and 1-30μm along the direction of the stress (depending on how the tip is modified, such as fluorescent dye, foil, etc). The main measurement error terms arise as the probe interfaces with the complex and heterogenous embryonic tissues. For example, as the deflection increases the deviation of contact angle to the tissue from vertical becomes more significant. Imperfections during foil preparation can add errors to the tissue contact area and increase tissue damage. In thicker tissues, probe tip tracking is more error-prone due to reduced contrast in the images, although this could be improved by modified tips such as those with glue and fluorescent dyes which create a trackable pattern (with trade-off of spatial resolution). These factors should be considered when estimating the stress measured or inflicted. For a detailed discussion of sources of errors and how to estimate/control them, see Methods section 4.4.

Using TiFM, a stress measurement against a deforming tissue takes 10-30min to reach stalling (when further deflection of the probe by the tissue becomes minimal, this may take longer when contact area is larger, such as in the case of a foil-probe construct) and thus the probe can be retracted shortly to minimize long-term invasiveness. The sharp tip creates little tissue damage (such as tearing) even with a strong load. These features are advantageous as the tissues are measured more closely to their native state with no large local deformation caused by implants such as gels or droplets. By applying a well-controlled stress close to the endogenous force of the tissue *in vivo*, downstream cellular responses such as gene expression changes can now be studied in more physiologically relevant ranges and at reduced experimental noises that are often difficult to achieve with mechanical perturbations. Combining TiFM loading with genetic probes would also allow the molecular force reporters to be calibrated (Kim *et al*., 2015). The requirement of bottom microscopy and therefore a thin flat sample like the early chicken embryo can be overcome with self-detecting probes or alternative deflection detectors such as an interferometer. Future work will aim at improving the automation and throughput of TiFM and expanding its applications to rheological measurements and other model systems. TiFM shows promise as a broadly useful method adding to the expanding toolbox (Campàs, 2016) for understanding the physical mechanisms of morphogenesis and quantitative engineering of development in small tissues.

## Materials and Methods

### Eggs and embryo preparation

Wild type chicken (*Gallus gallus*) eggs were supplied by Charles River Laboratories and Medeggs Inc. Tg(CAG-GFP) (McGrew *et al*., 2008) chicken eggs were provided by Clemson university (originally by University of Edinburgh). Eggs were kept in monitored 15°C fridge for storage and 37.5°C ~60% humidity egg incubators for incubation. HH stage 10-12 embryos were used. The early-stage embryos are used under a tissue protocol and do not require an animal protocol per institution guidelines. To obtain the embryos for TiFM measurements, eggs were incubated for ~40 hours before opening for the EC culture (Chapman *et al*., 2001). The EC culture uses 2cm x 2cm pieces of filter paper (Whatman) with two adjacent 0.5cm holes in the center. Eggs were opened into a petri dish and the thick albumen on the top that covers the embryo and the vitelline membrane is swept aside gently with small filter paper pieces using a tweezer. The holed filter paper is then lowered to attach to the vitelline membrane where embryos are visible through the hole (body axis aligned to the long axis of the hole). The vitelline membrane is then cut around the filter paper to release the embryo. The filter culture embryo is then rinsed in PBS to remove excess yolk. The cleaned embryo is then placed on a 3.5cm petri dish containing 2ml of culture gel made with the following formula (per 100ml of culture gel): Part A: 50ml Albumin (beaten for 15min) then supplement with 0.8ml 20% D-Glucose (Sigma); Part B: 0.3g BactoAgar (Sigma) solved in 50ml water in a microwave then supplement with 1.23ml 5M NaCl. Warm part A and cool part B to 55°C in a water bath. Mix thoroughly and add to petri dishes (2ml each) before gelation. The embryo cultures are then stored in a slide box with wet paper towels in the incubator. In experiments where some tissue areas and cells are labelled by DiI, the DiI was injected with a sharp-tipped glass needle by mouth pipetting from the ventral side of the embryo. The stock solution of 2.5mg/ml DiI in ethanol was diluted in PBS to 0.5mg/ml before injection. At the sample loading step of the TiFM procedure (see below), two embryos are taken and transferred to a pre-warmed glass bottom imaging dish (MatTek) covered with 200μl of culture gel. A second piece of filter paper is then added to prevent the embryo from detaching and floating once it’s submerged. Pre-warmed PBS was then added to cover the embryos, followed by 1-2 drops of mineral oil just enough to spread and cover the surface. One embryo is subjected to TiFM measurements/loading while the other serves as a control inside the incubation chamber on the scope.

### Design and Operation of the TiFM

A working TiFM can be assembled with the list of required equipment and components below. Design considerations are described and the components used in this study are listed, but it is not necessary to acquire the same components. The construction of the probe holder and incubation chamber would depend on the configuration of the base microscope that’s used. Similarly, existing microscope software can be incorporated into the operation procedure. Users with electrical engineering and programming experiences are required for the assembly and maintenance of the system.

#### 1. Required equipment and components

##### 1.1 Microscope

To construct a TiFM system, an inverted microscope with XY stage control and Z focus control is required. We used the Zeiss Axio Observer base (top modules including the TL illumination and condenser were removed). A low magnification objective 2.5-5x is required for sample positioning and a 10x objective is required for image data streaming.

##### 1.2 Camera

Due to the lack of TL illumination and the size and close proximity of the probe holder to the sample, side LEDs were included to compensate for the lack of light on the sample. A sensitive, fast camera is required to provide high resolution streaming of the probe tip in the tissue, which is essential for real-time feedback control of the force. The system’s temporal and spatial resolution limit is set by the camera and imaging protocol (described in more detail in section 4). We used a Ximea USB camera (MQ042MG-CM).

##### 1.3 Custom probe holder and capacitors

A stable, controllable probe needs to be installed on a reliable micromanipulator/stage as a holder with minimal drift over time. We used a World Precision Instruments WPI M3301R Manual Micromanipulator and a Newport Corp. 9062-XYZ-M stage. The stage was fixed to optical rails and beams (Thorlabs) onto the microscope base, forming an overhang on top of the sample stage. We 3D-printed 2-part plastic holders where the sample side has a slot for the chip of AFM probes and the piezo side has a slot for the insertion of the mobile end of the piezo (Fig. S2), and two slots on either side for the mobile copper capacitor plates. The 2 parts are tightened with screws to allow piezo and probe exchanges. The static end of the piezo is inserted in another printed holder which connects to a cage that holds the fixed capacitor plates. The capacitors flank the piezo whose positioning affects the capacitance difference (Figs. 1B-C), allowing a calibration of capacitance to holder position at the beginning of an experiment (described in more detail in section 4).

##### 1.4 Voltage controller and piezo

A programmable voltage controller is required to drive the piezo. We used a custom built one integrating a low and high power source but commercial ones such as Thorlabs (MDT694B) would also work. The controller needs to be able to adjust voltage output quickly and accurately during live measurements to enable the feedback control. The piezos we used are the ceramic piezoelectric benders from Thorlabs (PB4NB2W). Note that components in 1.3 should be designed in accordance with the type of the piezo and working range required.

##### 1.5 AFM probes

We used smooth (no tip modifications) silicon nitrate AFM probes from Bruker (MLCT-O10) and NanoAndMore (AIO-AL-TL) with spring constants ranging from 0.01-0.2N/m. Comparable probes will be feasible to use. It is advantageous to use probes with a larger length as they can reach deeper points of the embryo.

##### 1.6 Environmental chamber

The embryo requires proper temperature and humidity to develop normally. Common lab and microscope facility environments have a low humidity and room temperature around 25°C. We used a layer of mineral oil to reduce evaporation but this is not the optimal method as long-term survival of the embryo is affected under this condition. Environment chambers where humidity can be maintained at a high level while the electronics still function would be desirable. Alternatively, oxygenating the culture media that submerges the embryo may also be effective. We used a custom laser-cut cardboard box to enclose the holder and the sample stage and heating fans integrated with temperature sensors to maintain 37.5°C. Commercially available environmental chambers would also work but customization (e.g. additional holes) are needed to allow the installation of the probe holder and the in/out wires.

#### 2. Optional components

##### 2.1 Microcontroller

Because multiple data streams (images, capacitance, voltage, temperature, etc) flow through the system, it is advantageous to organize them under an integrated controller to align data onto the same time axis. For force measurement and loading small time differences in data streams do not cause a major issue because the probe and sample move slowly and errors average out through the feedback over time. However, for other measurements potentially capable by TiFM, such as oscillatory rheology, time axis alignment is critical. In these situations, a master clock is used to trigger the camera and capacitance readings for synchronization. We used a Teensy microcontroller to link different parts of the system and interface with Matlab (Mathworks) on the computer.

##### 2.2 Illumination modules

Having good contrast on the probe tip and embryo tissues is important for the precision of tip positioning and position measurement via image segmentation. The overhanging holder including the capacitors and the piezo form an occlusion for overhead TL illumination. Small LEDs can be installed on the holder to directly illuminate the sample. We used side LEDs which are fixed on the sample stage. The LEDs can be triggered by the camera or the microscope. We also used the fluorescence module of the Zeiss scope to provide RL illumination on the GFP transgenic embryos, and the probe tip which can be coloured by Quantum dots glued on via epoxy, mixing dyes such as DiI with the epoxy, or simply taking advantage of the autofluorescence of epoxy.

#### 3. Software

To measure and control forces, the locations of the chip and the tip are streamed in real time, and the chip location can be altered with piezo movement. Therefore, the main functionalities to achieve with the software are to send and receive the chip/piezo position (i.e. voltage), to program imaging, receive images and obtain tip position. We used Matlab (MathWorks) to create the user interface that plots the serial data via a USB link to the microcontroller, and to run the image segmentation. For the objective and shutter control we used Micromanager (Edelstein *et al*., 2014). A Teensy program that integrates the data streams is uploaded to the microcontroller prior to the start of experiments.

#### 4. Operation

##### 4.1 Testing and calibration

An assembled TiFM system needs to be tested and calibrated prior to loading actual samples. This step ensures the system functions properly and produces necessary parameters and data for the sample measurements. Firstly, holder stability must be tested for the desired duration of the experiment. Without samples, use the microscope to take a timelapse of the overhang holder without (a) and with the voltage controller on (b), and with the capacitor-voltage controller feedback loop on where a set capacitance is sent by the software (c). For a stable system, all timelapse results should show minimal movement of the probe, but the system is usable if (a) is stable and (c) achieves correction for drifts in (b). (a) tests the stability of the scaffold such as the rails, columns and the micromanipulator. (b) tests the stability of the piezo and voltage controller output. (c) tests the capacitor positioning system and the feedback. Secondly, the correlation between capacitance differences and the position of the holder needs to be established by driving the piezo across its dynamics range while capturing holder movement with timelapse imaging. This data serves as a lookup table that links capacitance reading to holder position, which will be used in sample measurements where the holder position can no longer be measured by imaging because of sample obstruction.

##### 4.2 Sample loading and probe insertion

Because the sample will result in light obstruction and scattering, the microscope view of the probe holder and tip will be blurry, making the probe insertion process difficult to see from the camera. Therefore, the XY position of the probe should be marked on the field of view prior to sample loading. The probe holder is then raised in Z (without touching its XY control) only to make room for the sample. After the sample dish (refer to “embryo preparation” for the protocol of readying a chicken embryo for TiFM) is in place, the desired tissue location can be aligned to the mark using the sample stage so that the probe tip will enter the right location once it’s lowered again. Once the probe enters the liquid layers, care must be taken to slowly lower the probe further to the desired tissue depth without overshooting which could cause tip breakage and/or tissue damage. The entrance of the probe/foil into the tissue is usually smooth because of their sharpness. Light conditions may make it difficult to see the location of the probe tip. It’s advisable to adjust the objective focus around the sample plane to find the probe tip. Once the probe is in the proper tissue location and in focus, camera and lighting settings can be further adjusted to ensure good-contrast images at a fast rate (low exposure time).

##### 4.3 Force measurement and loading

To measure tissue forces/stresses, the probe should be inserted to block the direction of tissue movement. If the desired measurement cross-section of the tissue is larger than the probe, a piece of aluminium foil can be attached to the probe tip via epoxy. We cut the foil pieces using a micromanipulator with a blade under a dissecting microscope to obtain rectangular pieces of 100-400μm. To determine the contact area between the tissue and the probe in order to calculate the stress (rather than just the total force), the insertion depth is measured by the Z positioning system of the microscope. Using the Zeiss Axio Observer as an example: first, the objective position is recorded from the Z-controller screen when the focus is on the surface of the tissue (e.g., endoderm for a dorsally mounted embryo such as in Fig. 1E); second, the objective is moved (lowered) to focus on the vicinity of the tissue layer of desired insertion depth (e.g., dorsal edge of the neural plate as in Fig. 1E); third, the probe (with or without foil) is inserted until the tip or edge of the foil is in focus at the desired depth, some minor adjustment of probe depth and/or focus might be performed for best focus and contrast, then the Z position of the objective is recorded again. Comparing the recorded objective Z positions yields the insertion depth. Using the insertion depth and known shapes of the probes and/or foils and the tissue, the tissue contact area can be estimated. To measure the stalling stress, once the tip/foil is in position, the capacitance should be fixed through the feedback loop to maintain the position of the holder. Timelapse imaging of the tissue area and the tip/foil displacement then indicates the force. The displacement will increase quickly then slowly and finally stall as morphogenesis is stalled by the probe. To load the tissue with a specific force, the force value will be evaluated against the current probe location and capacitance reading, and an adjustment of capacitance target (therefore holder position) will be sent to the voltage controller. With continued imaging and segmentation on the fly, the feedback loop maintains a dynamically stable difference between the holder and the probe tip (therefore cantilever deflection and force). After completion of measurements, the probe should be washed in deionized water by dipping to prevent damage from culture gel, albumen and salt crystals after drying.

##### 4.4 Sources and considerations of measurement errors

The cantilever (with force constant k) method’s working principle requires accurate measurements of the location of the holder (X_C_) and the tip (X_T_). For dynamical measurement and feedback control, these two measurements should also be synchronized in time. Therefore, factors that introduce inaccuracy for positional measurements and synchronization will bring error terms to the force measurements. In addition, because tissue stress (σ) is biologically more meaningful to calculate than the detected/inflicted force (F) which varies with tissue contact area (A), the requirement of contact area estimation raises additional error terms (σ=k(X_T_-X_C_)/A). For this current version of the TiFM, the largest error term is associated with the lowest resolution aspect of the system, which is the tracking of the tip position (hardware limited to ~200 frame/second and ~0.5μm pixel sizes). The capacitors and piezos are subject to fluctuations in the hardware such as from the voltage controller, but the sampling rate is higher and the errors are on a much smaller order of magnitude. Moreover, during force control, while there is a delay between imaging and segmentation of the tip to the action of voltage adjustment which may cause a force error, such errors will average out quickly over time through the feedback. Therefore the main error considerations focus on the spatial accuracy of X_T_ and the estimation of A. Segmentation and tracking of the tip (X_T_) without embryo sample (e.g., in air or water) produce high accuracy to the camera resolution. Errors increase as the imaging depth through tissue (D_I_) increases which deteriorates the tip image contrast/signal-to-noise ratio. This can be mimicked by imaging the probe movement behind increasingly thicker gels that scatter the light from the tip. In the case of the embryo the scattering will be additionally complex due to tissue heterogeneity. The exact positional uncertainty of the tip imaged through thick tissues depends on the conditions under which the images are obtained and should be taken into consideration when designing experiments. Using fluorescently labelled probes and surgically removing some tissues to image through are both effective ways of controlling this error. For the estimation of A, the insertion angle (θ_I_), depth (D) and features of the probe tip (e.g., dye, foil) should be considered. Taking the axis elongation force for example, the posterior body axis growth is largely horizontal during the stages concerned therefore a vertical insertion of probe is desirable. The insertion angle (θ_I_) is usually not perfectly vertical but can be adjusted by rotation of the mounting arm while moving the focal plane along the probe length (L) between the probe tip and base to minimize the on-camera horizontal movement. This can reach a sin(θ_I_)<0.05 for a L=200μm probe. The accuracy of depth of insertion (D) as obtained from the protocol described in 4.3 depends on the recognition of focal planes of tissue surfaces and probe tips by the user, and can in practice have ±20μm uncertainties which lead to uncertainties in contact area estimation. Depending on the type of tip, foils can have 10-20% uncertainty in A while narrow probes can only be accurate in the order of magnitude in terms of stress estimation under an error range of ±20μm in Z (e.g., Fig. 2C). Other factors include the quality of the foil surface and edges, where some curvature may make the effective A smaller than that of a flat foil. Effective ways in controlling the errors for A include: higher precision manufacturing of foils or other thinner materials (such as mica); transgenic fluorescent embryos which enhances the recognition of tissue layers/surfaces through focusing on cell layers. As an example, a well-preadjusted probe (θ_I_<15°) and a thin sample tissue location (such as the pPSM where both D_I_ and D<100μm) enables stress measurements by TiFM with a maximum 20% uncertainty term with a foil-probe construct (100μm wide), giving a high degree of confidence in the quantitative characterization of tissue forces.

## Data analysis

Movies were analyzed in Fiji (ImageJ). Probe/foil displacement was measured by object tracking in the intensity-time plot. The fluorescence intensities (as a proxy to cell density) were measured by drawing a ROI before and after the foil. DiI labelled cells were tracked with the Manual Tracking plugin. Tracking results and measurements were processed in Matlab (Mathworks) with custom scripts and plotted with Excel (Microsoft).

## Author contributions

C.U.C. and F.X. conceived the method with inputs from A.M, O.P. and L.M.; C.U.C. designed and constructed the TiFM with F.X.; A.M. contributed to hardware components and the oil incubation protocol. C.U.C., F.X. and J.M.N.V. performed the experiments. F.X. analysed the data and wrote the manuscript. The authors declare no competing financial interests.

## Acknowledgements

We thank S. Gapon, A. Mongera, and C. Guillot for logistical and technical assistance. This work is supported by an A*STAR International Scholarship to C.U.C., a NIH K99 award (HD092582) and a Wellcome Trust / Royal Society Fellowship (215439/Z/19/Z) to F.X., and a NIH R01 (HD097068) to O.P. and L.M.

## Data Availability

All data are presented in the paper. Source movies and custom codes are available upon request to

