## Supplemental information for "Direct Force Measurement and Loading on Developing Tissues in Intact Avian Embryos"

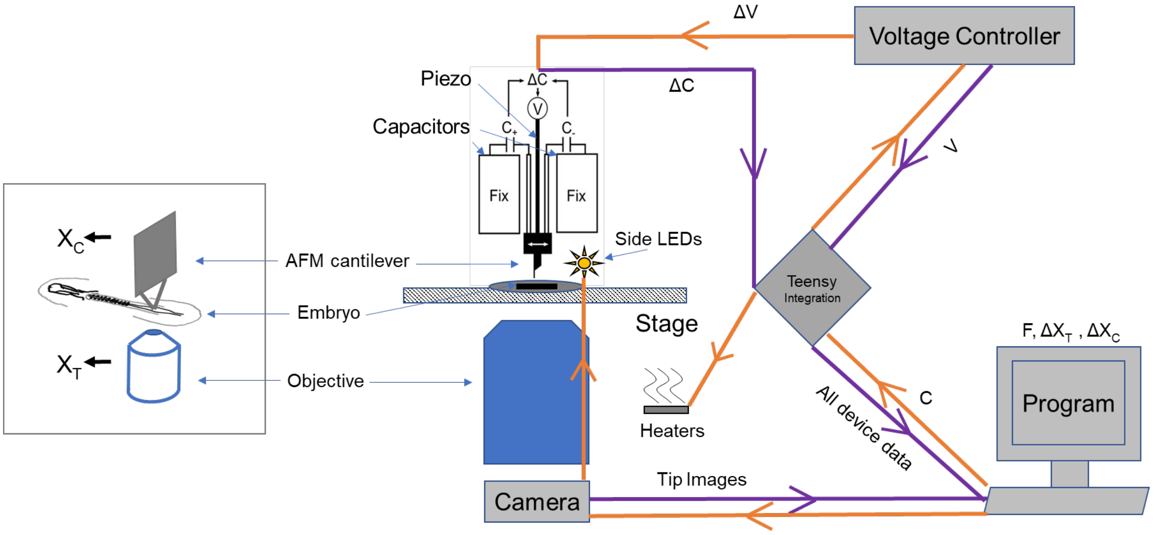


**Figure S1. Diagram of the TiFM system**
The left image is the conceptual design related to the actual system on the right. Purple arrows indicate data flow and orange arrows indicate command signals.

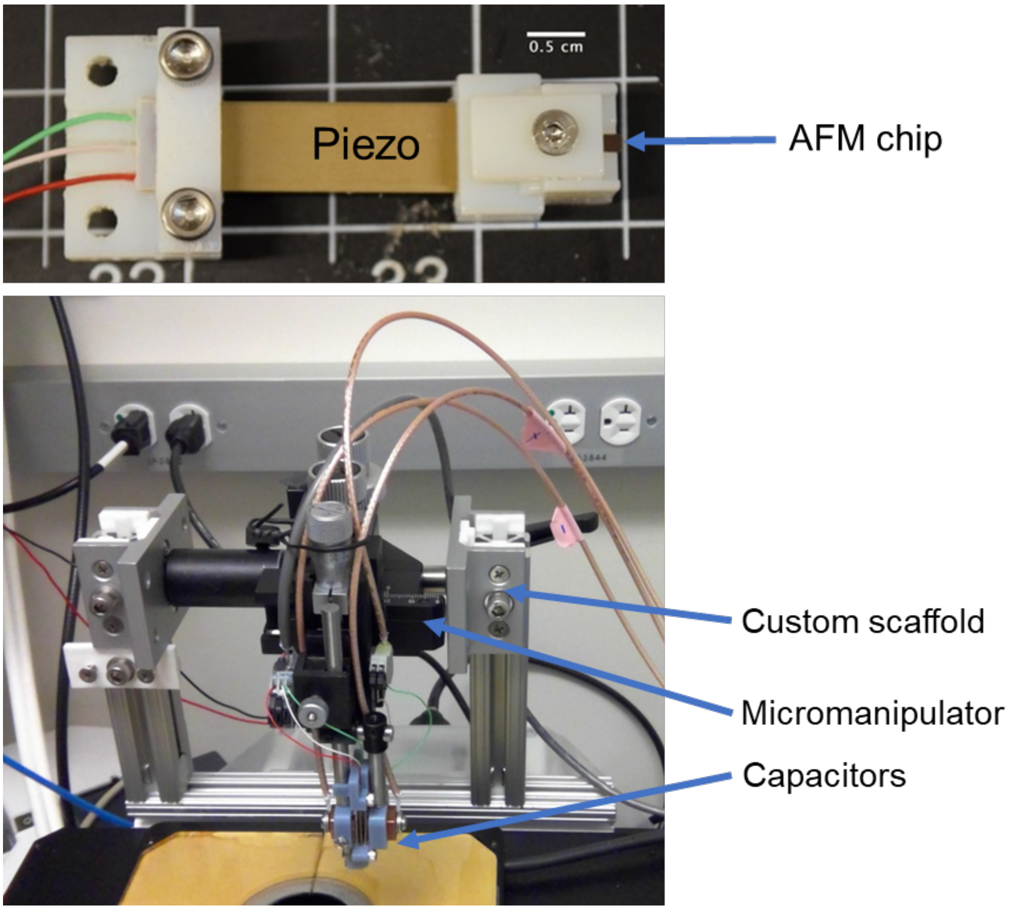


**Figure S2. Custom printed/built parts of the TiFM system**
Top image shows the 3D-printed plastic piezo holders. Bottom image shows the overhang scaffold. These designs are flexible with the actual scope and piezo used.

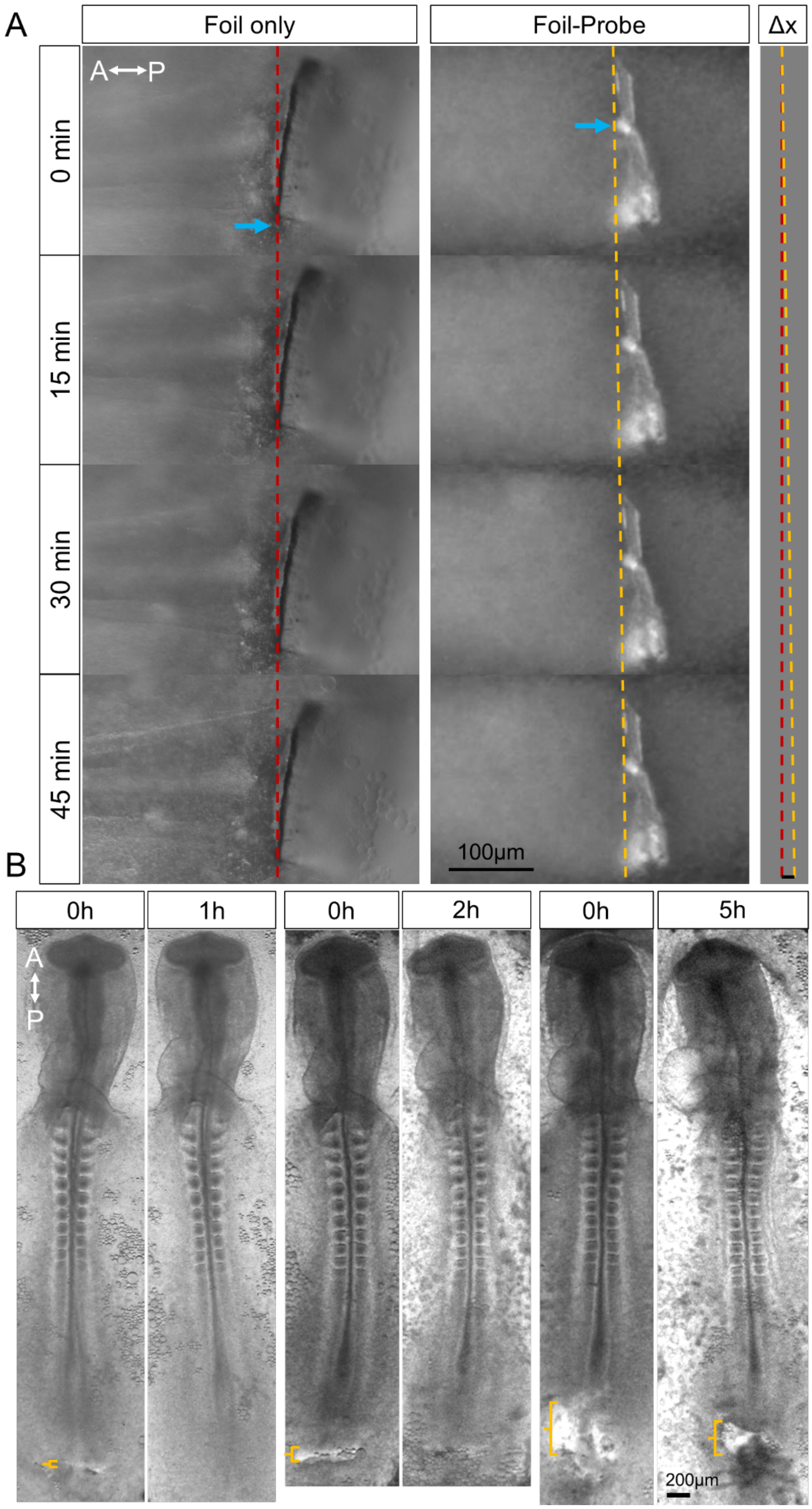


**Figure S3. Controls for foil insertion experiments**
(A) Tissue forces are insufficient to bend aluminium foils. A stripe of aluminium foil tailored to a ~400µm (L) x ~200µm (W) stripe was glued directly to the holder chip

rather than the cantilever probe. Tracking of this foil (blue arrows indicate a high-contrast point of the foil that was tracked) shows that it remains static, while the foil attached to the probe (from Fig. 1E) shows a displacement (Δx) of ~17μm. The force constant of the foils (k) can be estimated using the following information: the thickness ~20μm (t), Young's modulus of aluminium ~70GPa (E), and the second moment of area of rectangular beams $I=Wt^3/12$. Here $k=3EI/L^3 \approx 500$ N/m in the direction of the force under a free-end load condition (the value would be higher for the actual condition). This is significantly higher than the cantilever probes used (<1 N/m) in the experiments.
(B) Tissue invasiveness of the foil experiments as evaluated by wound healing. The end of the body axis was actuated with a foil to create different sized wounds. For a slit-like wound the tissue heals under 1hr, as the wound widens, healing becomes slower and/or incomplete over a longer duration when the wounds are too large. For stress measurements, the wounds are slit-like and heal quickly. Yellow brackets mark the width of the wound after foil extraction along the AP axis.

**Movie S1. Measuring axis elongation stress with TiFM**
Time stamps are hh:mm. This is a dorsal view of the tail end of a GFP embryo, anterior to the left. A small displacement of the foil is visible, providing information for inferring the stress. Refer to text and Fig. S3 for further details.

**Movie S2. Foil-only control for axis elongation stress with TiFM**
Time stamps are mm:ss. This is a dorsal view of the tail end of a WT embryo, anterior to the left. No stable displacement of the foil is observed (small short-term vibrations may be observed due to environmental noise). Refer to text and Fig. S3 for further details.

**Movie S3. Force loading on a live embryo**
Time stamps are hh:mm. Two GFP embryos, a control (left) and a loaded one (right) are shown, anterior to the top. Probe tip (modified with epoxy) is visible as a bright sphere with a dark bar in the center in the loaded embryo. The loaded embryo elongates at a faster rate. Refer to text for further details.

**Movie S4. Patterned tissue deformation on a loaded embryo**
Time stamps are hh:mm. This GFP embryo was loaded at 200nN and then unloaded. Anterior to the top. Probe tip is visible at the tail-end midline as a dark dot. Refer to text and Fig. 3A for further details.
